# Diosmetin alleviates liver inflammation by improving liver sinusoidal endothelial cell dysfunction

**DOI:** 10.1101/2023.10.23.563468

**Authors:** Dariusz Żurawek, Natalia Pydyn, Piotr Major, Krzysztof Szade, Katarzyna Trzos, Edyta Kuś, Ewelina Pośpiech, Piotr Małczak, Dorota Radkowiak, Andrzej Budzyński, Stefan Chłopicki, Jolanta Jura, Jerzy Kotlinowski

## Abstract

**Background & Aims:** Tumor necrosis factor-alpha (TNFα) induces pro-inflammatory activation in liver sinusoidal endothelial cells (LSEC) and liver inflammation. However, knowledge about whether modulating LSEC activation can alleviate liver inflammation is scarce. This study aimed to establish and validate an animal model mimicking LSEC dysfunction observed in patients with elevated plasma levels of TNFα, and explore whether vasoactive flavonoid diosmetin could serve as a therapeutic agent for liver inflammation.

**Approach & Results:** Genetic deletion of Mcpip1 in myeloid leukocytes (Mcpip1^fl/fl^LysM^Cre^) resulted in the development of systemic and liver inflammation in mice. Symptoms were compared with those in liver samples from obese humans with elevated TNFα. Mice were treated with diosmetin, and its effectiveness in alleviating liver inflammation was evaluated. Elevated TNFα correlated with reduced Mcpip1 expression in peripheral blood mononuclear cells and LSEC dysfunction in obese patients. Mcpip1 knockout in myeloid cells in mice replicated molecular signs observed in human samples. Diosmetin efficiently reduced LSEC activation and liver inflammation in Mcpip1^fl/fl^LysM^Cre^ mice. Diosmetin’s effects may stem from inhibiting NF-κB-p50 subunit production in TNFα-activated endothelial cells.

**Conclusions:** Diosmetin treatment efficiently restricted liver inflammation, despite ongoing systemic inflammation, by diminishing LSEC dysfunction. Mcpip1^fl/fl^LysM^Cre^ mice mimic symptoms of liver inflammation observed in humans and can be useful in studies on new anti-inflammatory therapies for the liver. We show that diosmetin, a vasoactive flavonoid that is successfully used in the clinic to treat chronic venous insufficiency, has also strong anti-inflammatory properties in the liver. This suggests that diosmetin treatment may be tested in humans as a supportive therapy for liver inflammation.

**Graphical abstract:** 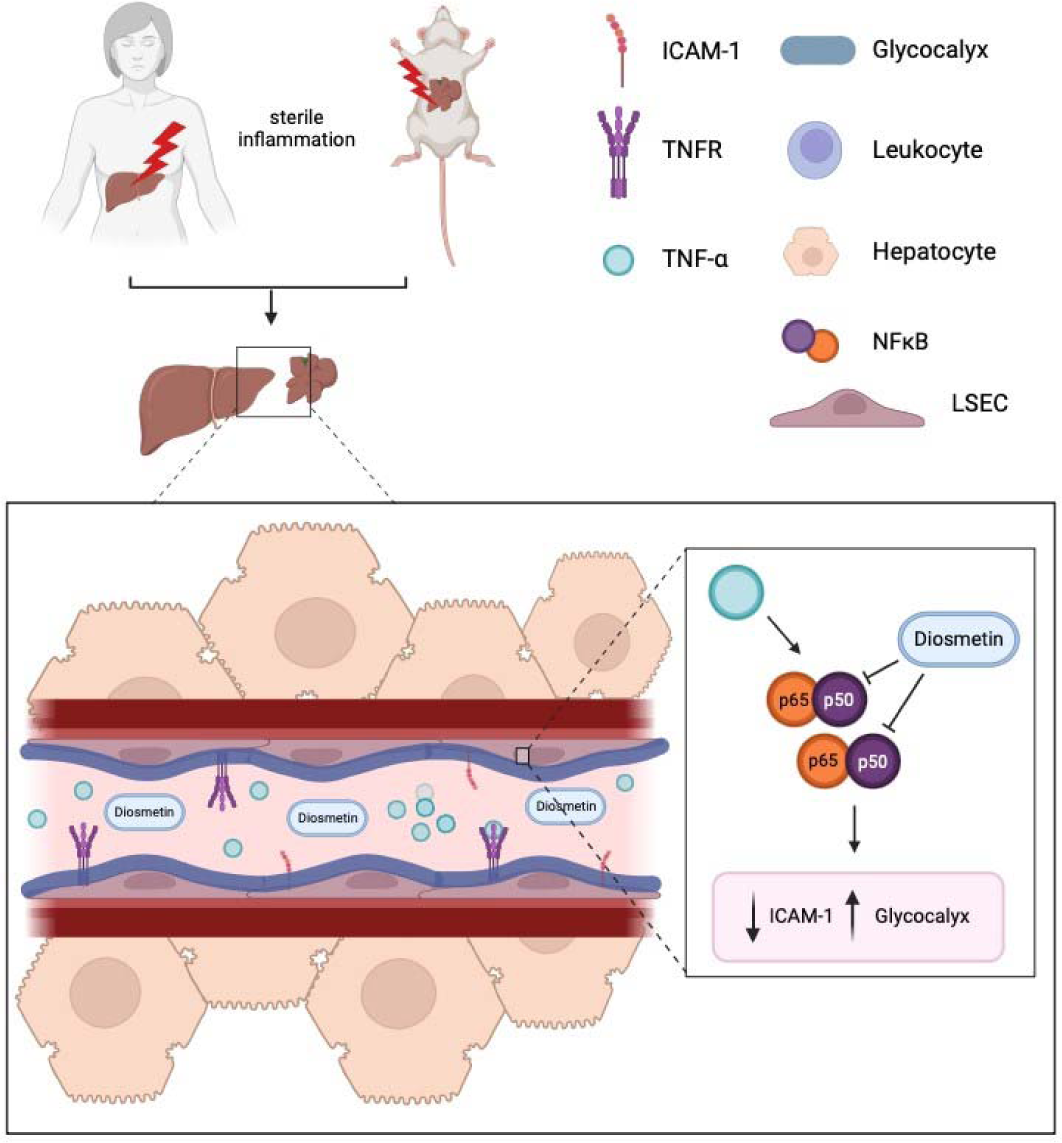

## Introduction

Sterile inflammation is expressed by the activation of the immune system despite the absence of infectious pathogens. It is a natural response of the body to tissue damage, and a necessary step in the wound healing process^1,2^. However, the sterile inflammatory response can become pathologically overactive and destructive for different types of tissues^1,2^. The liver is particularly prone to the damaging effects of prolonged or exacerbated sterile inflammation, which plays a central role in many liver diseases^3^. Genetic predispositions^4^, drug and toxin-induced liver injury^5^, and obesity associated with a diet rich in fatty acids^6^ are the main factors leading to the development of sterile liver inflammation. Recently, population-based studies reported that incidence rates of obesity-related nonalcoholic fatty liver disease (NAFLD)^7,8^, steatohepatitis (NASH)^8^ and alcoholic liver disease^9^ are estimated to be the leading causes of liver-related morbidity and mortality worldwide. Therefore, understanding the mechanisms underlying sterile liver inflammation and how to control them, is of prime importance.

Tumor necrosis factor alpha (TNFα) is a pro-inflammatory cytokine, produced by leukocytes of the myeloid lineage and T cells^10^, which plays a key role in the development of sterile liver inflammation. TNFα initiates and perpetuates liver inflammation in humans^11,12^ and animals^12,13^ by activating liver sinusoidal endothelial cells (LSEC)^14,15^, which in turn begin to produce on their surface cell adhesion molecules (CAMs)^16,17^. High levels of CAMs in the liver endothelium promote leukocyte-endothelial cell adhesion^18^, extravasation^19^ and, in effect, liver inflammation. Studies have shown that elevated blood levels of TNFα and LSEC dysfunction^20^ accompany multiple liver conditions, including autoimmune hepatitis (AIH)^21^ or NAFLD^22^ and NASH^11^ observed in obese patients. As a result, neutralization of TNFα have been proposed as a therapeutic strategy against AIH^21^, while bariatric surgery together with a lifestyle change are used as a treatment for morbid obesity and associated NAFLD, and NASH^23^. Although both therapeutic approaches are effective in alleviating the general symptoms of liver inflammation, they are associated with side effects. TNFα-targeted therapies increase susceptibility to infections and can cause autoimmune processes that develop *de novo* after their use^24^. Bariatric surgery, in turn, is an invasive surgical intervention that carries a risk of postoperative complications including acute liver failure^25,26^ or persistent liver inflammation^27^. Therefore, there is a need to develop adjuvant therapies that could help control liver inflammation and decrease the incidence rates of complications of currently used therapies. TNFα-dependent LSEC dysfunction is one of the key components of liver inflammation^28^; however, the knowledge of whether modulation of LSEC activation may be effective in preventing liver inflammation is sparse. Answering this question became the primary aim of the presented work.

The *Zc3h12a* gene encodes the Mcpip1 protein, a well-described endonuclease that negatively controls the levels of different pro-inflammatory cytokines, including TNFα^29,30^. The role of Mpcip-1 is to fine-tune inflammation and inhibit the development of exacerbated immune responses at systemic and organ-specific levels^29,31^. The Mcpip1 modifications are used as a genetic tool to generate an inflammatory phenotype in animal models.

Diosmetin (5,7-dihydroxy-2-(3-hydroxy-4-methoxyphenyl)chlormen-4-one) is a vasoactive flavonoid of polyphenolic structure^32^ and is widely found in fruits, vegetables, and certain beverages^33^. Diosmetin has clinically proven therapeutic effects in the treatment of chronic venous insufficiency^34^ and hemorrhoids^35^ by diminishing inflammatory activation of the endothelium and improving its function^36^. However, the usefulness of diosmetin as a supportive therapy for liver inflammation has not been studied. Therefore, testing whether anti-inflammatory modulation of LSEC by diosmetin may be efficient in alleviating liver inflammation observed in Mcpip1^fl/fl^LysM^Cre^ mouse model became a complementary objective of the presented work.

## Materials and Methods

### Generation of mice with *Zc3h12a* knockout in myeloid cells

Mice with loxP sites flanking exon 3 of the *Zc3h12a* gene were crossed with the mouse strain having nuclear-localized Cre recombinase inserted into the first ATG of the lysozyme 2 gene (LysMCre^Tg/Wt^, Stock#: 004781, Jackson’s Laboratory, USA). Mcpip1^fl/fl^ females were crossed with Mcpip1^fl/fl^ LysMCre^Tg/+^ males to obtain wild-type Mcpip1 ^fl/fl^ mice and Mcpip1^fl/fl^ LysMCre^Tg/Wt^ knockout mice (Mcpip1^fl/fl^LysM^Cre^). All mice had C57BL/6N background, were housed in ventilated cages under specific pathogen-free conditions in a temperature-controlled environment with a 14/10_h light/dark cycle, with food and water access *ad libitum*. For genotyping, DNA was extracted from tail tissue using the KAPA Mouse Genotyping Kit (KAPA Biosystems, USA) according to the manufacturer’s instructions. Genotyping for loxP insertion was performed by PCR (Supplementary file 1).

### Diosmetin treatment

3-month-old Mcpip1^fl/fl^ and Mcpip1^fl/fl^LysM^Cre^ mice were administered daily and *per orale* with 40mg/kg^37^ of diosmetin (Merck Milipore, Germany) dissolved in 200 ul of saline with 10% DMSO or vehicle. The diosmetin formulation was administered to mice with the use of a 10G oral gavage needle. Treatment lasted for the next three month until the animals reached 6 months of age. All *in vivo* experiments were conducted in accordance with the Guide for the Care and Use of Laboratory Animals (Directive 2010/63/EU of the European Parliament) and were carried out under the license no. 255/2018 issued by the 2^nd^ Local Bioethics Committee in Kraków.

### Mouse plasma and liver collection

Experimental mice were terminally anesthetized with a mixture of ketamine (150 mg/kg) and xylazine (35 mg/kg). Blood was collected by cardiac puncture in EDTA-coated tubes and centrifuged at 1500xg for 20 min. Plasma samples were collected and stored at −80°C. Whole livers were harvested, immersed in 10% buffered formalin solution (Merck Milipore, Germany) and kept for 48 h at 4 ° C.

### Human blood, peripheral blood mononuclear cells (PBMC), and liver sampling

Blood and liver samples were collected from 42 patients (10 men and 32 women) in The Second Department of General Surgery at Jagiellonian University Medical College (Kraków, Poland). Exclusion criteria included HCV/HBV/HIV infection, autoimmune diseases, cancer, and alcohol abuse. Of the 42 patients, 15 were diabetic. Blood was collected from fasting patients prior to bariatric surgery. All blood tests were routinely measured on the day of hospital admission in university hospital laboratories with ISO 9001 certificates, using comparable laboratory methods. To obtain PBMC, blood was collected in EDTA-coated tubes, diluted with PBS 1:1, layered on Lymphoprep™ (STEMCELL technologies, Canada) density gradient medium, and centrifuged for 20 mins at 800 xg. PBMC were harvested from appropriate density layer according to manufacturer’s protocol. Liver biopsies were taken during bariatric surgery and fixed in 10% buffered formalin solution. All human tissue samples were collected after obtaining written consent from patients according to the established protocol approved by the Local Bioethical Committee (study no. 122.6120.263.2016).

### Screening of inflammatory biomarkers in plasma

The mouse pre-mixed multianalyte kit (R&D Systems, Inc., USA) containing analyte-specific antibodies attached onto magnetic microparticles embedded with fluorophores was used to screen a set of 46 mouse biomarkers in plasma according to the manufacturer’s protocol. Plasma samples were analyzed in duplicates on a MAGPIX platform (Luminex, USA). The median fluorescence intensity (MFI) and a weighted 5-parameter logistic curve fitting method were used to calculate analyte concentrations.

### Human plasma TNF**α** measurement

DuoSet ELISA (R&D Systems, USA) was used to test the concentrations of TNFα in human plasma samples according to the manufacturer’s protocol. Absorbance was measured at 450 nm using Tecan Spectra Fluor Plus microplate reader (Tecan, Switzerland).

### Leukocyte analysis in mouse liver

Livers were perfused with 15 ml of Krebs buffer and cut into small pieces. Tissue fragments were digested in 10 ml of digestion medium (RPMI, 100U of penicillin, 0.1 mg/mL streptomycin, 1 mg/mL collagenase IV, 20 µg/mL DNAse I) at 37 ° C for 45 minutes under agitation (220 rpm) and filtered through a 100 µm cell strainer. The erythrocytes were lyzed in 5 mL of RBC lysis buffer (5 min, RT), the remaining cells were washed with 5 ml of PBS, filtered through a 40 µm cell strainer and centrifuged (300g, 4°C, 10 min), resuspended in 90 ul of PBS containing 2% FBS and incubated with 1 µL anti-mouse CD16/CD32 antibodies at 4 °C for 10 min. Leukocytes were stained (4°C, 30 min) with appropriate antibodies (Supplementary file 1) and DAPI. Cell acquisition was performed using a BD LSRFortessa (BD Biosciences, USA) flow cytometer and samples were analyzed using BD FACSDiva 8.0 software.

### Histological analysis of mouse and human livers

Fixed mouse and human liver samples were processed using the standard paraffin embedding method^38^, cut into 10 μm sections using a microtome (Leica Biosystems, Germany) and placed on microscope slides (Fisher Scientific, CA, USA). Tissue sections were deparaffinized with standard protocol^39^, followed by heat-mediated epitope recovery (30 min, 100 °C) in anhydrous citric acid solution (1.92 g of anhydrous citric acid per 1000 ml of distilled water, pH 2). Liver sections were stained with hematoxylin and eosin (H&E) according to standardized procedure^40^.

#### Glycocalyx

Liver sections were blocked for 30 min at RT in PBS with 5% BSA followed by three washes in PBS for 3 min. Tissue sections were then incubated for 60 min at RT with wheat germ agglutinin (WGA) conjugated with fluorescein isothiocyanate or rhodamine (ThermoFisher, USA) and diluted 1:1000 in PBS with 0.1% BSA. All sections were washed three times for 3 min with PBS and mounted with Dako Fluorescent Mounting Medium (Agilent, USA).

##### Mouse liver adhesion molecules

Autofluorescence was quenched, by incubation of tissue slices in copper(II) sulfate in acetate buffer (0.8 g CuSO_4_ dissolved in 500 ml of 50 mM ammonium acetate, pH 5) for 60 min at RT followed by two 2 min washes in ddH_2_O. Tissues were blocked for 60 min at RT in PBS containing 10% goat serum and 0.05% Triton X100. Slices were incubated overnight at 4 °C with appropriate primary antibodies in PBS with 0.1% Tween20 (PBS-T) and 5% BSA (Table S4). Nex, tissue sections were washed 3 times for 5 min in PBS-T, incubated for 60 min in RT with appropriate secondary antibodies (Fig. S1), washed, and mounted with Dako Fluorescent Mounting Medium (Agilent, USA).

#### Human liver ICAM1

The EnVision G2 double stain system, rabbit / mouse (DAB + / Permanent red) (Agilent, CA, USA) along with anti-human ICAM-1 primary antibodies (1:100) were used, according to the manufacturer’s protocol. Liver samples were counterstained with Mayer’s hematoxylin (Merck Milipore, Germany) and mounted with Glycergel Mounting Medium (Agilent, USA).

Images were taken with the Leica DMI6000B inverted wide-field fluorescence microscope with the 10× objective with Leica LAS X image acquisition software. All images were taken under the same illumination parameters, time of acquisition, and gain. For a quantitative analysis of the signal intensity, we analyzed 2-3 images of the liver per mouse and 5 images per human subject. Images were transformed to 8-bit files and analyzed using ImageGauge software (Fujifilm, Japan).

### LSEC isolation

LSEC were isolated from 6-month-old Mcpip1^fl/fl^ and Mcpip1^fl/fl^LysM^Cre^ mice according to a previously published protocol^41^. Briefly, livers were perfused with perfusion buffer (37°C) followed by perfusion with buffered collagenase (Liberase TM (Hoffman La-Roche, Switzerland). Parenchymal cells were removed using low-speed centrifugation at 50xg for 2 min. LSEC were isolated by gradient density sedimentation of resting non-parenchymal cells on 25–50% Percoll (Merck Millipore, Germany) and purified by using CD146-based isolation on magnetic MicroBeads (MACS, MiltenyiBiotec, Germany).

### RNA-seq analysis

Total RNA was extracted from LSEC by using mirVana miRNA Isolation Kit (Life Technologies, USA) according to the manufacturer’s protocol. Integrity and concentrations of the RNA samples were determined using an Agilent 2100 bioanalyzer with RNA 6000 Nano Kit (Agilent, USA). Libraries for 6 samples were prepared using the Ion Total RNA-Seq Kit v2 according to the manufacturer’s protocol using 12-22 polyA-enriched RNA as input material. Emulsion PCR, templating, and chip loading were performed manually. Libraries were barcoded, pooled in equimolar concentrations and sequenced on the Ion ProtonTM machine using Ion PI Hi-Q Sequencing 200 chemistry. Primary bioinformatic analyzes including quality check, read mapping to mm10 (with an average of 26 million reads mapped / sample) and transcript count were carried out using Torrent Suite Server v5.4.0 and RNA Seq Analysis v5.4.0.1 plugin. Normalization and analysis of differentially expressed genes (DEGs) were carried out using the DESeq2 package (with default parameters) implemented in the R version 3.3.3 software. The p values for differentially expressed genes were corrected for multiple comparisons using the Benjamini-Hochberg approach, and the results with corrected p-values <_0.05 were considered significant.

### Bioinformatic analysis

The gene ontology (GO) enrichment analysis of DEGs in LSEC was performed by using the GeneCodis 4 web tool (https://genecodis.genyo.es). GO biological processes were determined using hypergeometrical distribution analysis; corrected p-values <0.05 were considered significant. The signaling pathway impact analysis, which combines information on the fold change of DEGs and the topology of pathways, was performed using Graphite Web (https://graphiteweb.bio.unipd.it/index.html); pathways with corrected p-values < 0.05 were considered significant. The gene interaction network related to TNFα marker was assembled automatically by INDRA (http://indra.bio) by processing the available biomedical literature with multiple machine reading systems and integrating the selected pathway databases with obtained DEGs.

### RT-PCR

200 ng of total RNA was reverse transcribed into cDNA using High-Capacity cDNA Reverse Transcription Kit (ThermoFisher, USA) according to the manufacturer’s protocol. The mixture for each RT-PCR reaction contained 1x RT HS-PCR Mix SYBR Master Mix (A&A Biotechnology, Poland), 100 nM of appropriate custom-designed forward and reverse primers (Table S2) and 1ul od 10x diluted cDNA. Real-time PCR reactions for each gene were run under standard cycling conditions and automatic Ct threshold values on the QuantStudioTM 3 real-time PCR System (ThermoFisher, USA).

### Cytokine stimulation of human umbilical vein endothelial cells (HUVEC)

HUVEC cells were seeded in a 12-well plate (1 x 10^5^ cells/well), cultured for 48 h at 37°C with 5% CO2 in EGM™-2 Endothelial Cell Growth Medium-2 BulletKit™ (Lonza, UK) with penicillin–streptomycin (100_U/ml, Lonza, UK) and stimulated for 24 h with three different concentrations of cytokines found to be elevated in 6-month-old Mcpip1^fl/fl^LysM^Cre^ mice (Supplementary file 1). Untreated HUVEC cells were used as controls for the experiment. Stimulated cells were rinsed 3x in PBS and lysed in 30 ul of RIPA buffer with protease and phosphatase inhibitors.

### Co-stimulation of HUVEC cells with TNF**α** and diosmetin

HUVEC cells, seeded at a density of 1 x 10^5^ cells/well in a 12-well plate, cultured for 48 h at 37°C with 5% CO2 in EGM™-2 Endothelial Cell Growth Medium-2 BulletKit™ (Lonza, UK) with penicillin–streptomycin (100_U/ml, Lonza, UK) were co-stimulated with a mixture of TNFα (8pg/ml or 10 ng/ml) and diosmetin (2uM, 20uM or 40uM) for 24h, washed 3x with PBS and lysed in 30ul of cold RIPA buffer with protease and phosphatase inhibitors. Control cells were left untreated.

### Immunoblotting

Immunoblotting was performed according to the standardized protocol^42^. 30 ug of total protein per sample was loaded onto an 8% polyacrylamide gel, separated and transferred to nitrocellulose blotting membranes (Merck Milipore, Germany). The membranes were incubated with appropriate primary and secondary antibodies (Supplementary file 1). Proteins were visualized using Lumi-Light Western Blotting Substrate (Roche, Switzerland). The signal was detected using the ChemiDOC MP Imaging System (Bio-rad, Germany). Signal density was analyzed with the use of ImageJ software.

### Statistical analysis of the results

The Shapiro-Wilk normality test was applied to check the Gaussian distributions of all data sets. Cytokine profiling in mouse plasma, mouse liver histochemistry, RT-PCR and HUVEC co-stimulation with TNFα and diosmetin were analyzed using Two-Way ANOVA with Benjamini, Kreiger, Yekutieli post hoc test correcting the results for multiple comparisons (FDR). Mouse liver cytometry, human and mouse liver histological quantification were analyzed using an unpaired t-test. HUVEC cell stimulation with different concentrations of cytokines was analyzed using one-way ANOVA followed by the post hoc FDR test. The Mann-Whitney test was used to compare human biochemical analysis. Spearman’s correlation was used to analyze the associations between plasma TNFα and liver ICAM1 and glycocalyx levels in humans and mice. Logistic regression and calculation of the ROC curve were used to estimate the predictive potential of ICAM1, glycocalyx, and combination of both on plasma TNFα levels in humans and mice. All results are expressed as mean ± SEM and were analyzed using either the GraphPad Prism9.0 or Statistica13 software.

## Results

### Decreased Mcpip1 levels in PBMC correlate with increased plasma levels of TNFα and LSEC dysfunction in patients

We measured TNFα concentrations in plasma samples from a cohort of 42 patients who had undergone bariatric surgery at the University Hospital in Krakow. Patients were divided into two experimental groups: controls with normal TNFα (i.e. below 14 pg/ml)^43^, and patients with elevated TNFα (i.e. above 14pg/ml)^43^ (Fig. 1A). Both groups were matched in terms of demographic and biochemical parameters (Supplementary table 1). We identified significantly decreased expression levels of Mcpip1 protein in PBMC of patients with high TNFα (Fig. 1B and 1D) compared to controls. Additionally, Mcpip1 levels in PBMC negatively correlated with plasma levels of TNFα (Fig. 1C). Histological analysis revealed a significantly higher expression of ICAM1 in LSEC of patients with elevated TNFα as compared to the control group (Fig. 1E upper panel and Fig. 1F). ICAM1 levels positively correlated with plasma TNFα concentrations (Fig. 1G). Furthermore, we observed a significant decrease in glycocalyx levels in LSEC of patients with high TNFα (Fig. 1e lower panel and Fig. 1h) as compared to controls. In this case, LSEC glycocalyx levels negatively correlated with TNFα concentrations in patients (Fig. 1I). Logistic regression analysis demonstrated that liver levels of ICAM1 and glycocalyx had strong potential to predict the TNFα-dependent systemic inflammatory phenotype in humans (Fig. 1J).

**Fig. 1.**
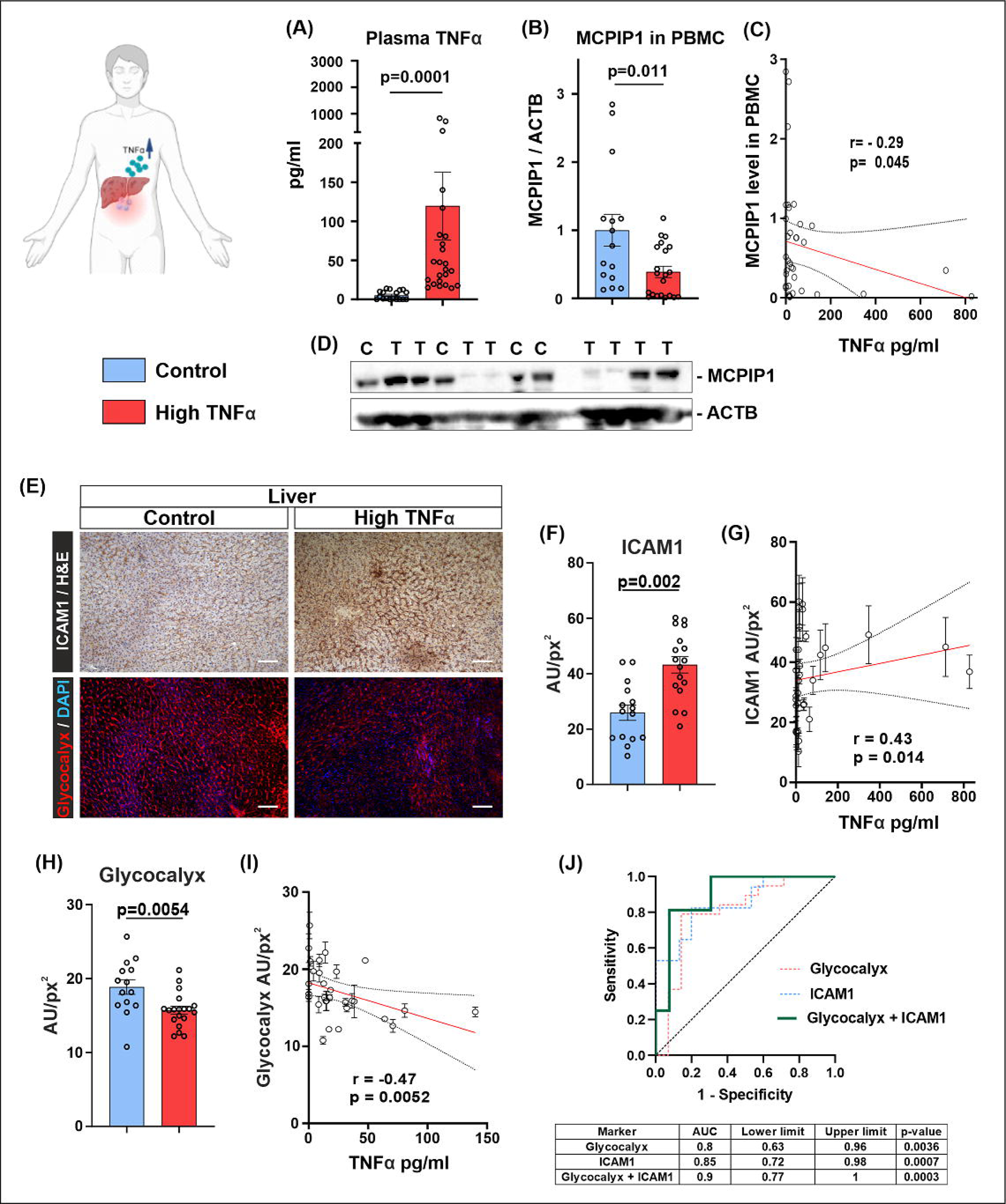
Patients with elevated plasma levels of TNFα exhibit reduced MCPIP1 expression in PBMC and signs of LSEC dysfunction. **(A)** TNFα concentration in human plasma (n=18-24, p=0.0001, Mann-Whitney test). (**B)** MCPIP1 protein levels in human PBMC (n=15-21, p=0.011, Mann-Whitney test). **(C)** Spearman’s correlation analysis between plasma TNFα concentrations and MCPIP1 levels in PBMC of patients. **(D)** Representative western blot of MCPIP1 and b-actin in PBMC of patients. C – control patients with normal TNFα, T – patients with high TNFα. **(E)** IHC pictures of ICAM-1 (upper panel) and Glycocalyx (lower panel) in LSEC of patients with normal and elevated TNFα, scale bar = 50 μm. **(F)** Densitometric quantification of ICAM-1 expression in human livers (n=15-17, p=0.002, unpaired t-test). **(G)** Spearman’s correlation analysis between plasma TNFα concentration and liver ICAM1 levels in patients. **(H)** Densitometric quantification of glycocalyx in human livers (n=14-19, p=0.0054, unpaired t-test). **(I)** Spearman’s correlation analysis between plasma TNFα concentration and liver glycocalyx levels in patients. **(J)** ROC curve (simple logistic regression) estimating the predictive potential of the liver levels of ICAM-1, glycocalyx, and combination of both on the presence of systemic inflammation reflected by plasma TNFα levels in patients. Data are represented as means ± SEM in the bar graphs. Abbreviations: ACTB – β-actin, AU/px^2^ – Auxiliary Units per squared pixel, H&E – Hematoxylin and Eosin staining, ICAM1 - Intercellular Adhesion Molecule 1, MCPIP1 – monocyte chemoattractant protein induced protein-1, PBMC – peripheral blood mononuclear cells, TNFα – tumor necrosis factor alpha.

### Mcpip1 knockout in myeloid cells leads to the development of systemic and liver inflammation in mice

We employed the Cre-loxP system to selectively delete Mcpip1 in myeloid cells (Fig. 2A) and generated the Mcpip1^fl/fl^LysM^Cre^ mouse strain to mimic observations made on human samples. Specificity of Mcpip1 knockout in Mcpip1^fl/fl^LysM^Cre^ mice was confirmed by its reduced expression in bone marrow, with no change in LSEC or liver samples compared to control mice (Fig. 2B). Mcpip1^fl/fl^LysM^Cre^ model is well-established, and its cell-type specificity has been extensively studied^44^. Deletion of Mcpip1 in myeloid cells induced systemic inflammation in Mcpip1^fl/fl^LysM^Cre^ mice, which progressed gradually over their lifespan, reflected by the altered plasma cytokine profiles (Fig. 2C). 6-month-old Mcpip1^fl/fl^LysM^Cre^ mice showed elevated TNFα, interleukin-6 receptor α, chemokines regulating myeloid cell function (MCP-2, MCP-5, MIP-1α, MIP-3α, MIP3β), and two soluble markers of endothelial cell activation and damage, i.e. CHI3-L1 and Syndecan-1 (Sdc1)^45,46^ compared to 3-month-old Mcpip1^fl/fl^LysM^Cre^ and control animals of both age groups (Fig. 2 D-E). We observed massive inflammatory infiltrates in the liver parenchyma of 6-month-old Mcpip1^fl/fl^LysM^Cre^ animals (Fig. 2F – lower panel), while 3-month-olds exhibited only sporadic foci of initial inflammation (Fig. 2F – upper panel). This observation suggests a close association between the development of liver inflammation and the presence of systemic inflammation in mice. Mcpip1 deletion in myeloid leukocytes (Cd11^+^ cells) led to the expansion of this population (Fig. 2G), with increased percentages of Ly6C^+^ and Ly6C^+^MHCII^+^ myeloid subpopulations, but not neutrophils (Cd45^+^Cd11b^+^Ly6G^+^), in the livers of 6-month-old Mcpip1^fl/fl^LysM^Cre^ mice as compared to controls (Fig. 2I). Furthermore, the livers of 6-month-old Mcpip1^fl/fl^LysM^Cre^ mice contained reduced amounts of B cells (Cd45^+^Cd11b^-^B220^+^) and significantly increased percentages of T cells (Cd45^+^Cd11b^-^Cd3^+^) as compared to controls (Fig. 2H).

**Fig. 2.**
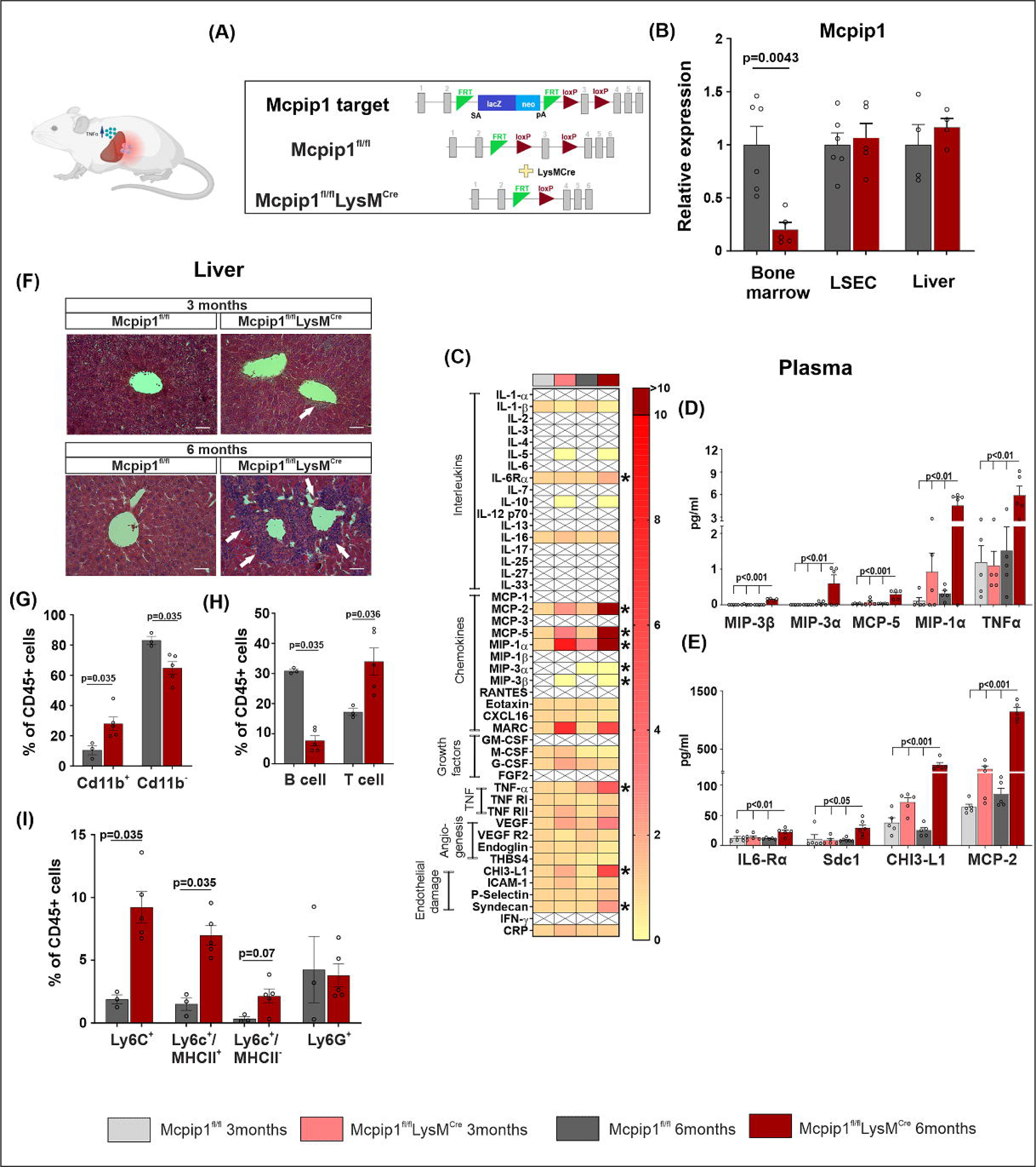
Deletion of Mcpip1 in myeloid cells leads to the development of systemic and liver inflammation in Mcpip1^fl/fl^LysM^Cre^ mouse model. **(A)** Schematic representation of Mcpip1 knock-out model. **(B)** Expression of mRNA encoding Mcpip1 in Bone marrow (n=5-6, p=0.0043, Mann-Whitney test), LSEC (n=5-6, p=ns, Mann-Whitney test), and Liver (n=5-6, p=ns, Mann-Whitney test) in Mcpip1^fl/fl^LysM^Cre^ and control mice. **(C)** Heatmap showing screening results of 46 markers in mouse plasma samples (n=4-6, * p<0.05 for two-way ANOVA followed by FDR post-hoc test), X – marker non detected. **(D-E)** Concentration levels of significantly changed plasma markers in mice (n=4-6, two-way ANOVA followed by FDR post-hoc test). **(F)** H&E staining images representing 3- and 6-month-old wild type (Mcpip1^fl/fl^) and Mcpip1 knock-out (Mcpip1^fl/fl^LysM^Cre^) mouse livers. White arrows point inflammatory infiltration, scale bars – 50 μm. **(G)** Cytometric analysis of myeloid Cd11b^+^ (n=3-5, p=0.035, Mann-Whitney test) and non-myeloid Cd11b^-^ (n=3-5, p=0.036, Mann-Whitney test) leukocytes in 6-month-old Mcpip1^fl/fl^LysM^Cre^ and control mouse livers. **(H)** Cytometric analysis of B-cells (n=3-5, p=0.035, Mann-Whitney test) and T-cells (n=3-5, p=0.036, Mann-Whitney test) in 6-month-old Mcpip1^fl/fl^LysM^Cre^ and control mouse livers. **(I)** Cytometric analysis of Ly6C^+^ (n=3-5, p=0.035, Mann-Whitney test), Ly6C^+^MHCII^+^ (n=3-5, p=0.035, Mann-Whitney test), Ly6C^+^MHCII^-^ (n=3-5, p=0.07, Mann-Whitney test) myeloid subpopulations together with Ly6G^+^ neutrophils (n=3-5, p=ns, Mann-Whitney test) in 6-month-old control and Mcpip1^fl/fl^LysM^Cre^ mouse livers. Data are presented as mean ± SEM in the bar graphs. Abbreviations: CHI3-L1 - Chitinase 3 Like 1, IL-6Rα – Interleukin-6 receptor-alpha, MCP-5 - Monocyte Chemotactic Protein-5, MCP-2 - Monocyte Chemotactic Protein-2, Mcpip1 – monocyte chemoattractant protein induced protein-1, Mcpip1^fl/fl^ – control mice, Mcpip1^fl/fl^LysM^Cre^ – mice with Mcpip1 knock-out in myeloid cells, MIP-1α - macrophage inflammatory protein-1-alpha, MIP-3α - macrophage inflammatory protein-3-alpha, MIP- 3β - macrophage inflammatory protein-3-beta, Sdc1 – Syndecan-1, TNFα – tumor necrosis factor alpha, p=ns – p-value non significant.

### Mcpip1 knockout in myeloid cells is associated with elevated plasma levels of TNF**α** and LSEC dysfunction in Mcpip1^fl/fl^LysM^Cre^ mice, mimicking similar changes observed in humans

RNA-seq analysis of LSEC from 6-month-old Mcpip1^fl/fl^LysM^Cre^ mice, exhibiting fully developed systemic and liver inflammation, revealed 1206 DEGs as compared to controls (Fig. 3A-B). These DEGs were most enriched in the cell adhesion pathway (Fig. 2E), the only significantly activated biological pathway in LSEC taking into consideration the directions of changes in DEGs (Fig. 3F). Gene network analysis pinpointed TNFα as an upstream signaling marker regulating the expression of DEGs enriched in the CAMs pathway (Fig. 3D). RT-PCR validation confirmed increased expression levels of CAMs and decreased expression of Sdc-1 in LSEC of 6-month-old Mcpip1^fl/fl^LysM^Cre^ mice compared to their 3-month-old littermates and control animals of both ages (Fig. 3C).

**Fig. 3.**
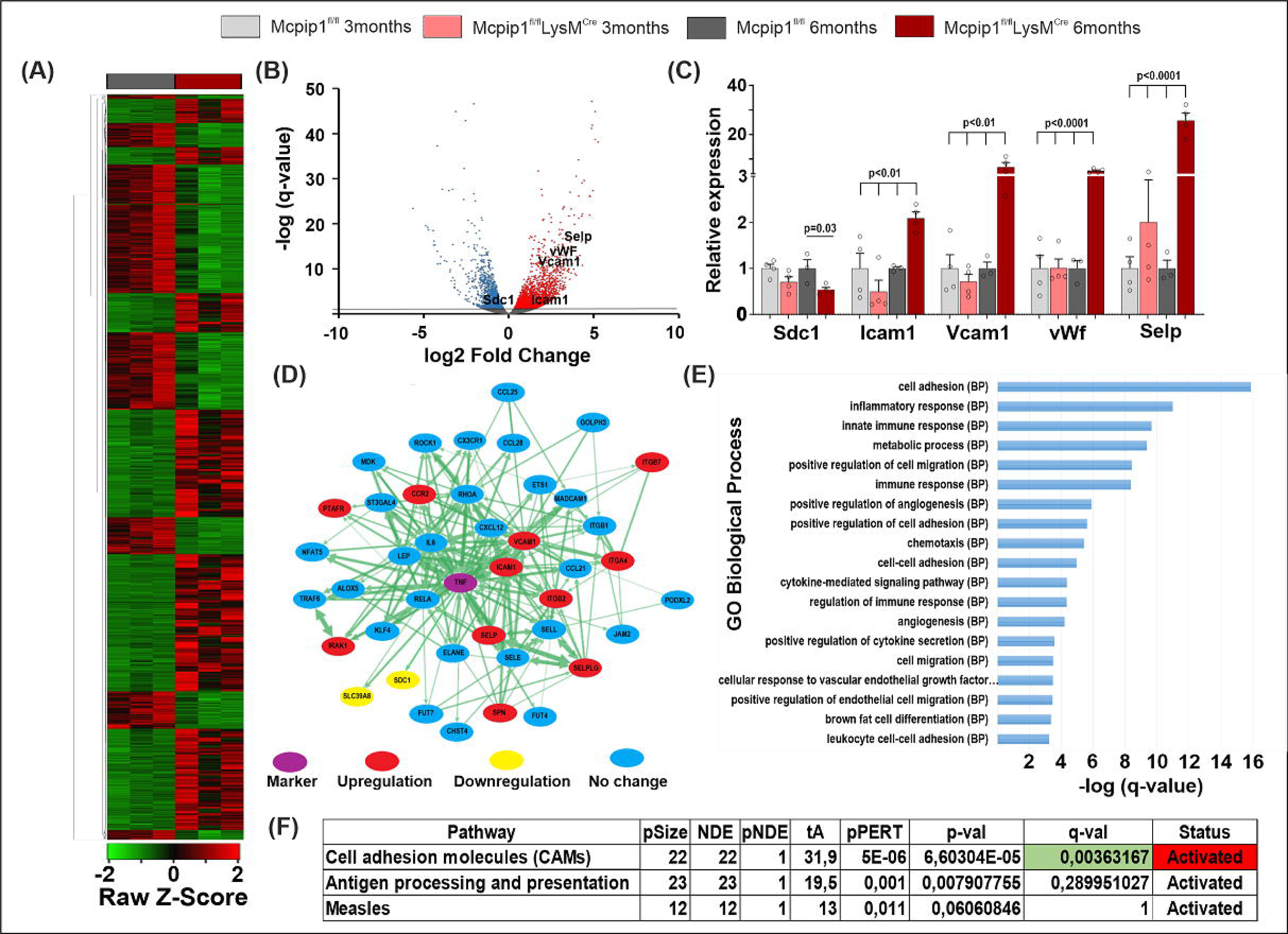
LSEC from Mcpip1^fl/fl^LysM^Cre^ mice with developed systemic inflammation exhibit transcriptomic changes characteristic to pro-inflammatory activation and cell dysfunction. **(A)** Heatmap and **(B)** volcano-plot representing DEGs in LSEC from 6-month-old control and Mcpip1^fl/fl^LysM^Cre^ mice (n=3). **(C)** Expression of mRNAs encoding Sdc1, Icam-1, Vcam-1, vWf and Selp genes in LSEC of 3- and 6-month-old control and Mcpip1^fl/fl^LysM^Cre^ mice (n=3-4, p-values calculated by two-way ANOVA followed by FDR post hoc test). **(D)** The gene interaction network analysis pointed out that TNFα is an upstream signaling marker for DEGs in LSEC from 6-month-old Mcpip1^fl/fl^LysM^Cre^ mice. **(E)** Gene ontology analysis of DEGs in LSEC of 6-month-old Mcpip1^fl/fl^LysM^Cre^ mice. **(F)** Signaling pathway impact analysis of DEGs showed that cell adhesion molecules is the only significantly activated biological pathway in LSEC from 6-month-old Mcpip1^fl/fl^LysM^Cre^ mice. Data are presented as mean ± SEM in the bar graphs. q-val <0.05 considered as significant. Abbreviations: Icam1 - Intercellular Adhesion Molecule 1, NDE – number of DEGs in the pathway, pNDE - hypergeometric probability, pPERT - bootstrap probability associated to tA,, p-Size - the number of genes in the pathway, p-val – nominal p-value, q-val – FDR adjusted p-value, Sdc1 – Syndecan-1, Selp – selectin P, STATUS – Inhibition/Activation according to the negative/positive sign of tA., tA- perturbation score value, Vcam1 Vascular cell adhesion molecule 1, vWF – von Willebrandt factor.

IHC analysis revealed significantly increased expression levels of Icam-1 (Fig. 4A-B), Vcam-1 (Fig. 4C-D) and Selp (Fig. 4E-F) in LSEC of 6-month-old Mcpip1^fl/fl^LysM^Cre^ mice as compared to controls and 3-month-old Mcpip1^fl/fl^LysM^Cre^ mice. Since Sdc-1 is the main transmembrane glycoprotein of endothelial glycocalyx and its downregulation is considered a marker of endothelial cell dysfunction^47^, we examined whether changes at the level of mRNA encoding Sdc1 may be followed by changes in surface glycocalyx (Glx) levels in LSEC. We found a significant Glx decrease in LSEC of 6-month-old Mcpip1^fl/fl^LysM^Cre^ mice as compared to control and young Mcpip1^fl/fl^LysM^Cre^ animals (Fig. 4G-H).

**Fig. 4.**
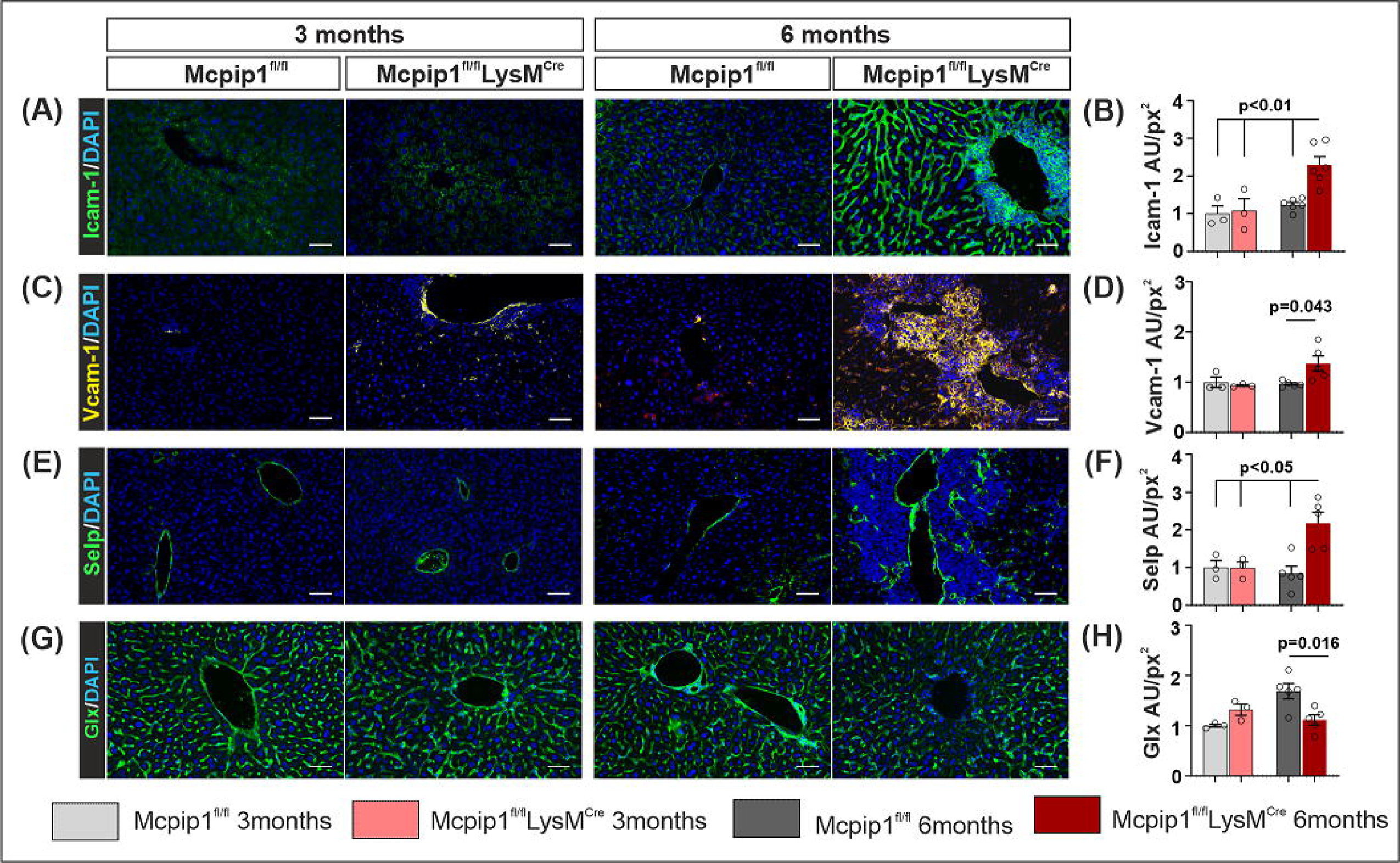
Time-dependent development of LSEC dysfunction and liver inflammation in Mcpip1^fl/fl^LysM^Cre^ mice with Mcpip1 deletion in myeloid cells. **(A)** IHC images of Icam1 levels in LSEC of 3- and 6-month-old control Mcpip1^fl/fl^LysM^Cre^ mice, scale bars – 50 μm. **(B)** Densitometric quantification of Icam-1 expression in mouse livers (n=3-6, p<0.01, two-way ANOVA followed by FDR post hoc test). **(C)** IHC images of Vcam1 levels in LSEC of 3- and 6-month-old control and Mcpip1^fl/fl^LysM^Cre^ mice, scale bars – 50 μm. **(D)** Densitometric quantification of Vcam-1 expression in mouse livers (n=3-6, p=0.043, two-way ANOVA followed by FDR post hoc test). **(E)** IHC images of Selp levels in livers of 3- and 6-month-old control and Mcpip1^fl/fl^LysM^Cre^ mice, scale bars – 50 μm. **(F)** Densitometric quantification of Selp expression in mouse livers (n=3-6, p<0.05, two-way ANOVA followed by FDR post hoc test). **(G)** IHC images of Glx levels in LSEC of 3- and 6-month-old control and Mcpip1^fl/fl^LysM^Cre^ mice, scale bars – 50 μm. **(H)** Densitometric quantification of Glx levels in mouse livers (n=3-6, p=0.016, two-way ANOVA followed by FDR post hoc test). Data are presented as mean ± SEM in the bar graphs. Abbreviations: Icam-1 - intercellular Adhesion Molecule 1, AU/px^2^ – arbitrary units per square pixel, Glx – glycocalyx; Mcpip^fl/fl^ – control mice, Mcpip1^fl/fl^LysM^Cre^ - mice with Mcpip1 deletion in myeloid cells, Selp – selectin P, Vcam-1 - vascular cell adhesion molecule 1.

Complementary *in vitro* experiments with HUVEC endothelial cells indicated that, among the elevated plasma cytokines in Mcpip1^fl/fl^LysM^Cre^ mice, only TNFα was able to stimulate CAMs (Fig. S1A-C and S1E-J) and reduce SDC1 expression levels (Fig. S1D and S1E-J). We utilized HUVEC cells as a well-established *in vitro* model for studying various endothelial cell functions, including liver vasculature^48^, under pro- inflammatory conditions^18^. Recent lineage tracing experiments support the idea that a substantial portion of liver vasculature originates from the sinus venosus endocardium, indicating a shared ancestry between HUVEC and LSEC^49^. Furthermore, HUVEC cells respond to TNFα through similar factors as LSEC^50^, including TNFα-dependent NF-κB activation^51^ and induction of ICAM1 expression^52^, establishing HUVEC as reliable surrogate endothelial cell line for studying LSEC responses to inflammation *in vitro*.

Additionally, we observed that 6-months-old Mcpip1^fl/fl^LysM^Cre^ mice exhibited multiple molecular features similar to those observed in our cohort of human patients. 6- months-old Mcpip1^fl/fl^LysM^Cre^ mice with elevated plasma levels of TNFα (Fig. 5A) and decreased levels of Mcpip1 in myeloid cells (Fig. 5B) showed increased expression levels of Icam1 (Fig. 5C upper panel and Fig. 5D) and decreased levels of Glx in LSEC (Fig. 5C lower panel and Fig. 5F). Similarly to human samples, Icam1 in LESC positively correlated with plasma TNFα levels (Fig. 5E), while LSEC Glx levels showed negative correlation with TNFα (Fig. 5G).). Logistic regression analysis demonstrated that liver levels of ICAM1 and glycocalyx had strong potential to predict the TNFα-dependent systemic inflammatory phenotype in mice (Fig. 5H), what was similar observation in humans.

**Fig. 5.**
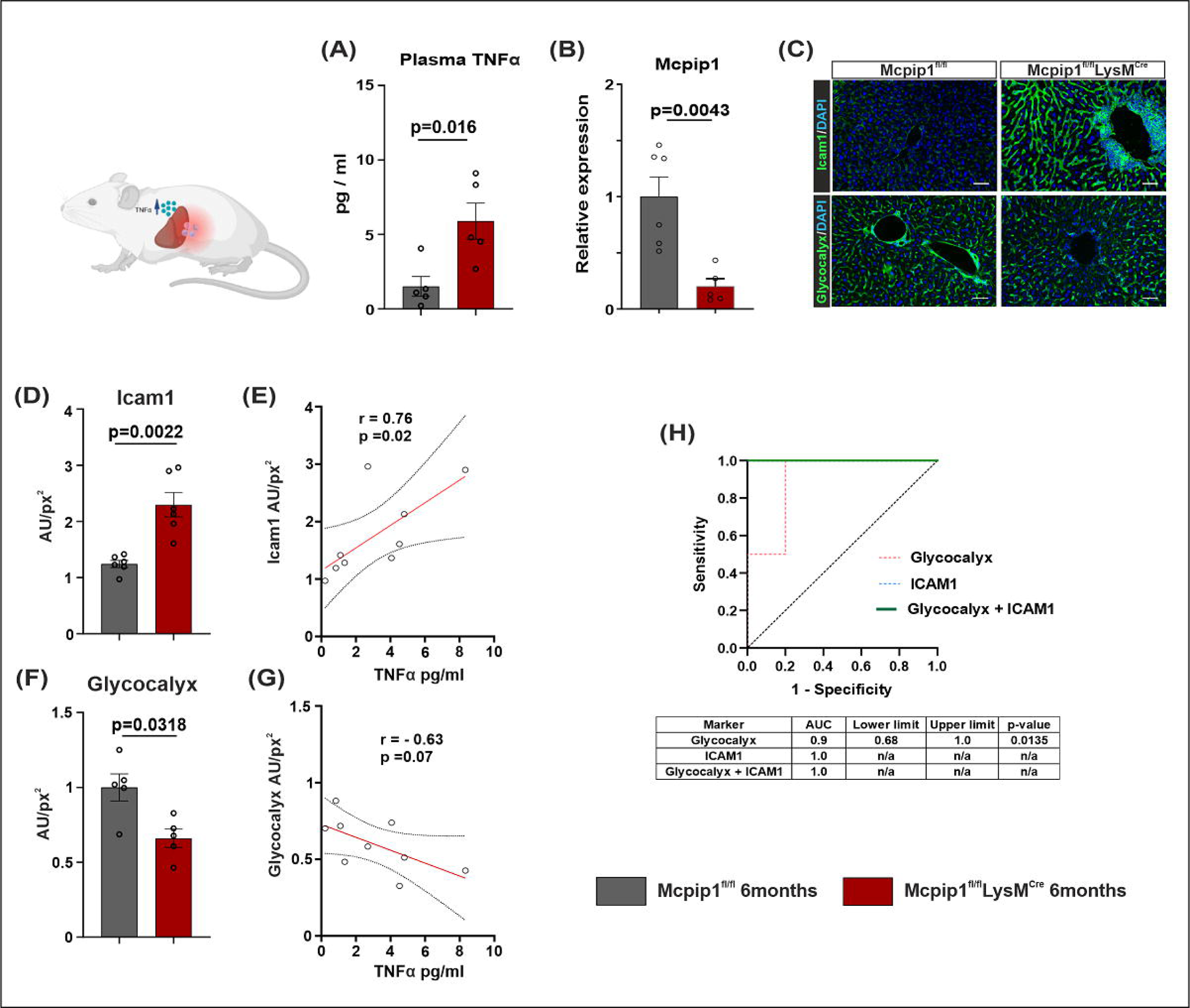
LSEC from 6-months-old Mcpip1^fl/fl^LysM^Cre^ mice exhibit molecular features similar to those observed in patients with high TNFα. **(A)** TNFα concentration in plasma of 6-months-old control and Mcpip1^fl/fl^LysM^Cre^ mice (n=5, p=0.016, Mann-Whitney test). **(B)** Expression of mRNA encoding Mcpip1 in bone marrow of 6-months-old control and Mcpip1^fl/fl^LysM^Cre^ mice (n=5-6, p=0.0043, Mann-Whitney test). **(C)** IHC images representing Icam1 (upper panel) and Glx (lower panel) levels in LSEC of 6-month-old control (Mcpip1^fl/fl^) and Mcpip1 knock-out (Mcpip1^fl/fl^LysM^Cre^) mice, scale bars – 50 μm. **(D)** Densitometric quantification of Icam-1 expression in 6-months-old control and Mcpip1^fl/fl^LysM^Cre^ mouse livers (n=6, p=0.0022, Mann-Whitney test). **(E)** Spearman’s correlation analysis between plasma TNFα concentration and liver Icam1 levels in 6-month-old mice. **(F)** Densitometric quantification of glycocalyx expression in 6-months-old control and Mcpip1^fl/fl^LysM^Cre^ mouse livers (n=5, p=0.0318, Mann-Whitney test). **(G)** Spearman’s correlation analysis between plasma TNFα concentration and liver glycocalyx levels in 6-month-old mice. **(H)** ROC curve (simple logistic regression) estimating the predictive potential of the liver levels of Icam1, glycocalyx, and combination of both on the presence of systemic inflammation reflected by plasma TNFα levels in 6-months-old control and Mcpip1^fl/fl^LysM^Cre^ mice. Data are presented as mean ± SEM in the bar graphs. Abbreviations: AU/px^2^ – arbitrary units per square pixel, Icam-1 - intercellular Adhesion Molecule 1, Mcpip1 – monocyte chemoattractant protein induced protein-1.

These findings highlight TNFα as the key cytokine activating LSEC in mice and humans and support our hypothesis that Mcpip1^fl/fl^LysM^Cre^ is well-validated animal model mimicking LSEC dysfunction observed in patients with elevated plasma levels of TNFα.

### Diosmetin treatment prevents liver inflammation in Mcpip1^fl/fl^LysM^Cre^ mice by decreasing the pro-inflammatory activation of LSEC

To assess whether modulation of LSEC activation by the vasoactive drug diosmetin can prevent liver inflammation *in vivo,* we treated 3-month-old Mcpip1^fl/fl^LysM^Cre^ mice with 40mg/kg of diosmetin, until they reached 6 month of age and developed an inflammatory phenotype (Fig. 6A). Diosmetin treatment effectively prevented liver inflammation in Mcpip1^fl/fl^LysM^Cre^ mice. Diosmetin-treated mice exhibited only small, single foci of leukocyte infiltration in the liver parenchyma, while untreated mice showed extensive leukocyte infiltrations (Fig. 6B). Importantly, diosmetin treatment did not affect the development of systemic inflammation (Fig. 6C). Plasma pro- inflammatory cytokine profiles, including TNFα, did not differ between untreated and diosmetin treated Mcpip1^fl/fl^LysM^Cre^ mice, but remained elevated as compared to control animals (Fig. 6D-E). However, plasma biomarker profiling revealed improved endothelial function expressed by a significant decrease in plasma Sdc1 levels in diosmetin-treated animals as compared to untreated littermates (Fig. 5E).

**Fig. 6.**
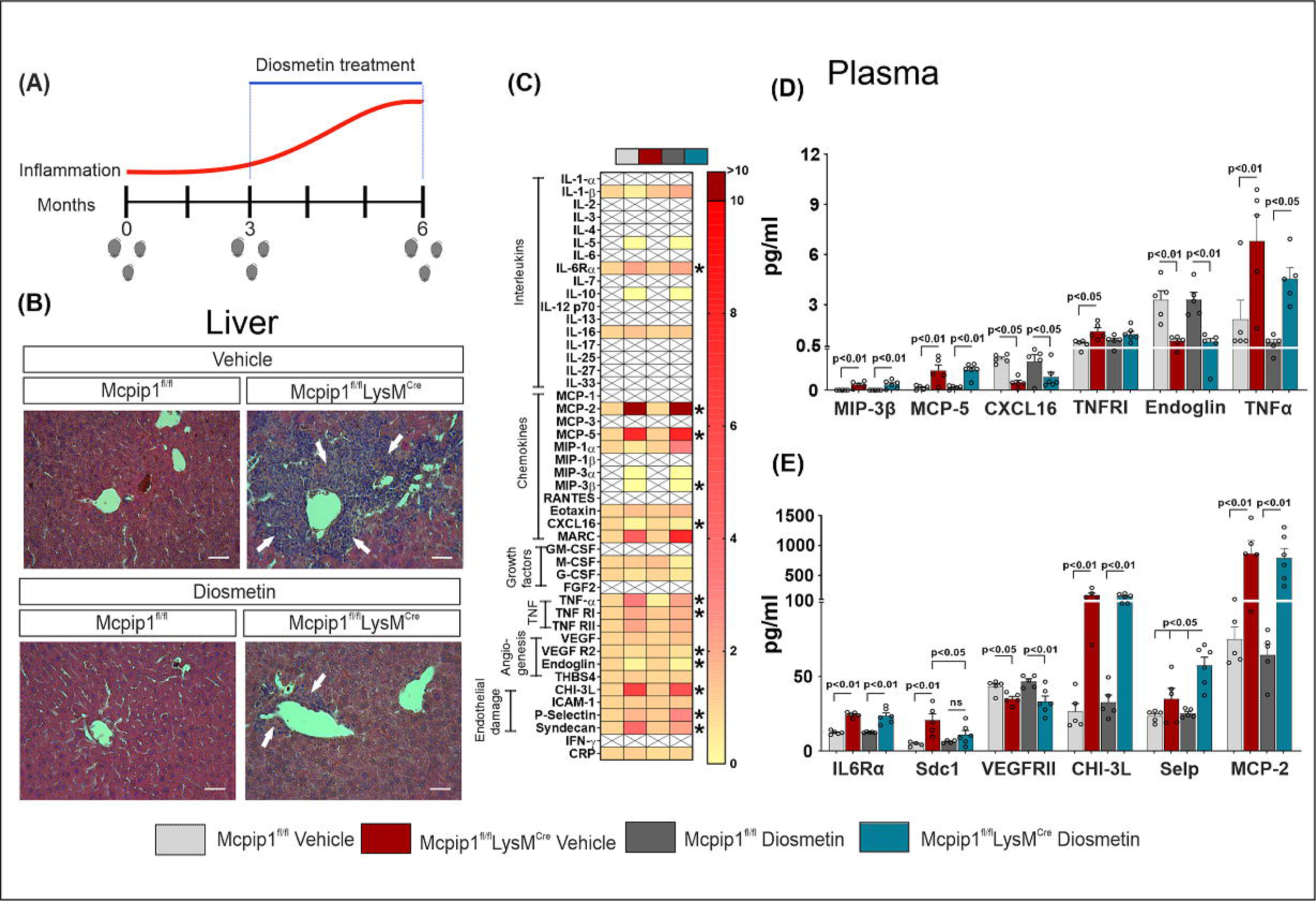
Diosmetin treatment prevents liver inflammation, but does not modulate the development of systemic inflammation and TNFα levels in Mcpip1^fl/fl^LysM^Cre^ mice. **(A)** Schematic representation of diosmetin treatment experiment *in vivo*; **(B)** H&E staining images representing livers from 6-month-old control and Mcpip1^fl/fl^LysM^Cre^ mice treated with diosmetin or vehicle for 3 months. White arrows point inflammatory infiltrations, scale bars – 50 μm; **(C)** Heatmap showing screening results of 46 markers in plasma samples from 6-month-old control and Mcpip1^fl/fl^LysM^Cre^ mice treated with diosmetin or vehicle for 3 months (n=4-6, * p<0.05, two-way ANOVA followed by FDR post-hoc test), X – marker non detected; **(D-E)** Concentrations of significantly changed plasma markers in 6-month-old control and Mcpip1^fl/fl^LysM^Cre^ mice treated with diosmetin or vehicle for 3 months (n=4-6, two-way ANOVA followed by FDR post-hoc test). Data are presented as mean ± SEM. Abbreviations: CHI3-L1 - Chitinase 3 Like 1, CXCL16 - Ligand chemokin 16, IL-6Rα – Interleukin-6 receptor-alpha, MCP-2 - Monocyte Chemotactic Protein-2, MCP-5 - Monocyte Chemotactic Protein-5, Mcpip1^fl/fl^ – wild type mice, Mcpip1^fl/fl^LysM^Cre^ – mice with Mcpip1 knock-out in myeloid cells, MIP-3β - macrophage inflammatory protein-3-beta, Selp – Selectin P, Sdc1 – Syndecan-1, TNFα – tumor necrosis factor alpha, TNFRI – TNFα receptor type I, VEGFRII - Vascular endothelial growth factor receptor 2.

Histological analysis confirmed that diosmetin treatment significantly reduced LSEC activation and improved its function. This was reflected by decreased expression levels of Icam-1 (Fig. 7A-B), Vcam-1 (Fig. 7C-D) and Selp (Fig. 7E-F), along with increased levels of glycocalyx (Fig. 7G-H) in LSEC of diosmetin-treated Mcpip1^fl/fl^LysM^Cre^ mice as compared to non-treated animals. Hence, the anti- inflammatory effect of diosmetin is related to the direct LSEC modulation rather than dampening systemic inflammation. This is supported by the observation that plasma TNFα levels (and other pro-inflammatory cytokines) remained significantly elevated in Mcpip1^fl/fl^LysM^Cre^ mice regardless of diosmetin treatment (Fig. 6C-E).

**Fig. 7.**
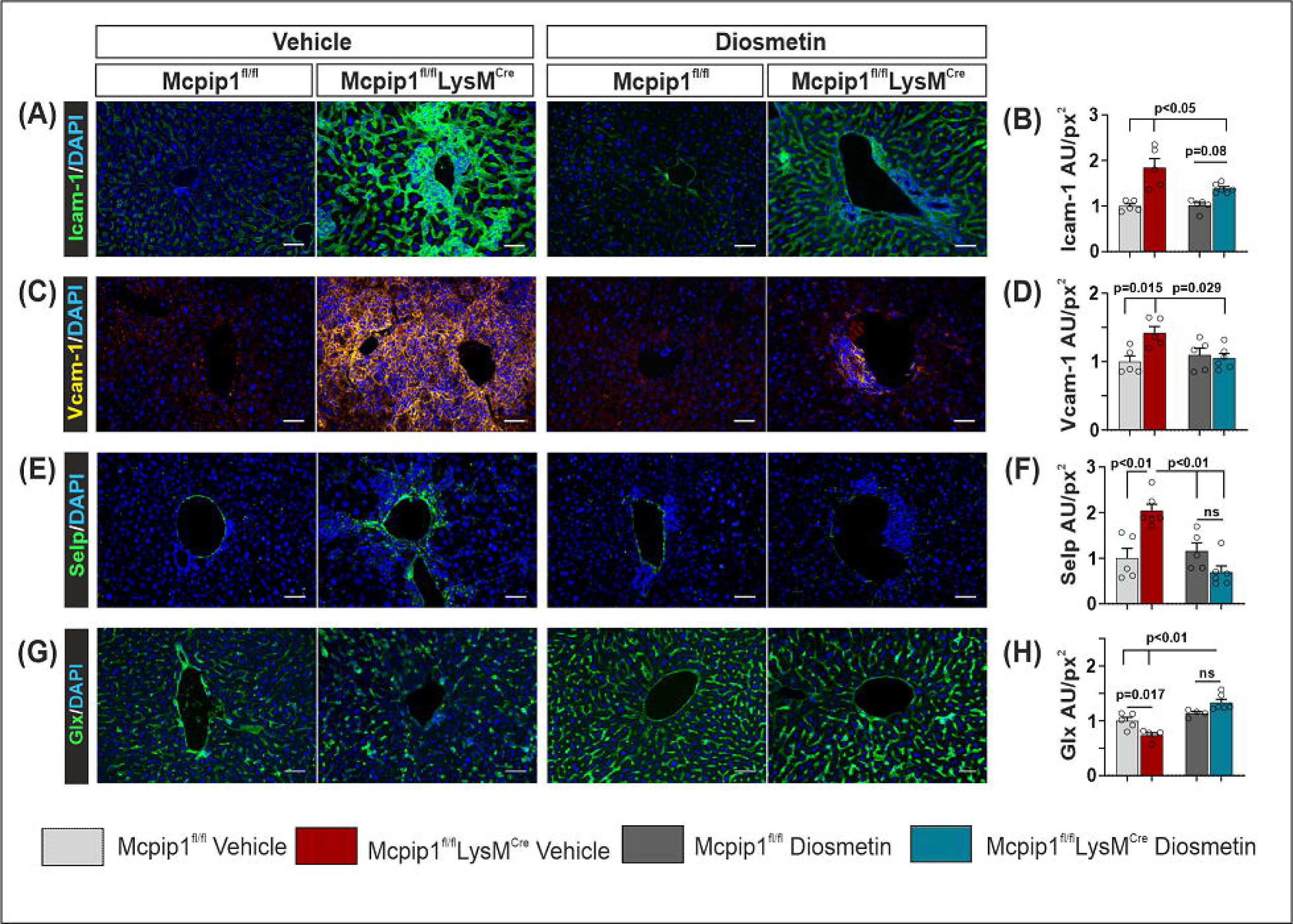
Diosmetin treatment significantly decreases pro-inflammatory activation and dysfunction of LSEC in Mcpip1^fl/fl^LysM^Cre^ mice. **(A)** IHC images of Icam1 levels in LSEC from 6-month-old control and Mcpip1^fl/fl^LysM^Cre^ mice treated with diosmetin or vehicle for 3 months. Scale bars – 50 μm. **(B)** Densitometric quantification of Icam-1 expression in livers of 6-month-old control and Mcpip1^fl/fl^LysM^Cre^ mice treated with diosmetin or vehicle for 3 months (n=5-6, p<0.05 and p=0.08, two-way ANOVA followed by FDR post hoc test). **(C)** IHC images of Vcam1 levels in LSEC from 6-month-old control and Mcpip1^fl/fl^LysM^Cre^ mice treated with diosmetin or vehicle for 3 months. Scale bars – 50 μm. **(D)** Densitometric quantification of Vcam- 1 expression in livers of 6-month-old control and Mcpip1^fl/fl^LysM^Cre^ mice treated with diosmetin or vehicle for 3 months (n=5-6, p<0.015 and p=0.029, two-way ANOVA followed by FDR post hoc test). **(E)** IHC images of Selp levels in livers from 6-month-old control and Mcpip1^fl/fl^LysM^Cre^ mice treated with diosmetin or vehicle for 3 months. Scale bars – 50 μm. **(F)** Densitometric quantification of Selp expression in livers of 6-month-old control and Mcpip1^fl/fl^LysM^Cre^ mice treated with diosmetin or vehicle for 3 months (n=5-6, p=0.01 and p<0.01, two-way ANOVA followed by FDR post hoc test). **(G)** IHC images of Glx levels in LSEC from 6-month-old control and Mcpip1^fl/fl^LysM^Cre^ mice treated with diosmetin or vehicle for 3 months. Scale bars – 50 μm. **(H)** Densitometric quantification of Glx expression in livers of 6-month-old control and Mcpip1^fl/fl^LysM^Cre^ mice treated with diosmetin or vehicle for 3 months (n=5-6, p=0.017 and p<0.01, two-way ANOVA followed by FDR post hoc test). Data are presented as mean ± SEM in the bar graphs. Abbreviations: AU/px^2^ – arbitrary units per square pixel, Icam-1 - intercellular Adhesion Molecule 1, Glx – glycocalyx, Mcpip^fl/fl^ – control mice, Mcpip1^fl/fl^LysM^Cre^ - mice with Mcpip1 deletion in myeloid cells, Selp – selectin P, Vcam-1 - vascular cell adhesion molecule 1.

Endothelial cell activation by TNFα, mediated by NF-κB transcription factor^53^, is a common mechanism observed in HUVEC^50^ and LSEC^54^. We found that diosmetin significantly blocked *in vitro* TNFα-induced increase in Icam-1 (Fig. S2A and S2H), Vcam-1 (Fig. S2B and S2H) and Selp (Fig. S2C and S2H) proteins in HUVEC. Diosmetin also prevented TNFα-induced SDC1 protein decrease (Fig. S2D and S2H). In vitro, diosmetin inhibited TNFα-induced generation of NFκB-p50 – a biologically active subunit of NF-κB transcription factor (Fig. S2G and S2I). Diosmetin did not interfere with neither TNFα-induced degradation of NF-κB inhibitory protein IκBα (Fig. S2E and S2I) nor NFκB-p105 subunit, the initial non-active form of NF-κB- p50 subunit (Fig. S2F and S2I). These results suggest that diosmetin is a potent drug that blocks the pro-inflammatory effect of TNFα primarily by inhibiting the generation of an active NFκB-p50 subunit in endothelial cells.

## Discussion

Liver inflammation, a common human condition, often leads to LSEC dysfunction, liver fibrosis, cirrhosis, and an increased risk of hepatocellular carcinoma. The global burden of liver diseases underscores the urgency of finding effective therapeutic interventions that will prevent liver inflammation. Our research revealed a significant correlations between elevated TNFα, decreased Mcpip1 protein levels in PBMC, and molecular symptoms of LSEC dysfunction in patients, suggesting that all three molecular factors may be engaged in common inflammatory-related mechanisms. To test this hypothesis *in vivo*, we induced Mcpip1 deletion in myeloid cells creating the Mcpip1^fl/fl^LysM^Cre^ mouse model. The Mcpip1 protein has established role in controlling the inflammatory state at both systemic and organ-specific levels^29,31,55^. Mcpip1 is an endonuclease that regulates inflammation by direct destabilization of mRNA transcripts encoding e.g. IL-1β or IL-6 cytokines^29,56^, and by indirect reduction of TNFα levels^30,56^. Importantly, the Mcpip1^fl/fl^LysM^Cre^ mouse model has previously been used to investigate the role of sterile inflammation in different organs^55–57^. Here, we show that Mcpip1^fl/fl^LysM^Cre^ mouse model effectively mimics sterile liver inflammation and LSEC dysfunction offering a tool for searching new anti- inflammatory therapeutic strategies for the liver.

Well-validated animal models of disease are important for scientific research, as they provide a reliable platform for testing potential treatments and understanding the pathomechanisms of different diseases. However, to be fully useful in biomedical research animal model must meet three basic validation criteria^58^. Face validation refers to the replication, by the animal model, of key symptoms or pathological changes observed in human disease. Predictive validation refers to the ability of the animal model to accurately predict human responses to treatments. Construct validation refers to the similarities of the underlying biological mechanisms of the animal model and human disease. Here, we show that Mcpip1^fl/fl^LysM^Cre^ mice meet all validation criteria and thus may serve as an animal model of sterile liver inflammation. Mcpip1 deletion in myeloid cells produced a systemic and liver inflammation characterized by increased TNFα, pro-inflammatory activation of LSEC, and expansion of monocytes and T lymphocytes into the liver parenchyma. The observed changes fulfill the face validity criterion, as they mimic the symptoms observed in our cohort of patients. Mcpip1^fl/fl^LysM^Cre^ mice showed key molecular aspects of inflammation and LSEC dysfunction found in humans i.e. reduction in the levels of Mcpip1 in myeloid cells, and an overexpression of CAMs and decreased glycocalyx levels mediated by TNFα in LSEC. Noteworthy is the remarkably strong correlation between ICAM-1 and glycocalyx expression levels in LSEC and plasma TNFα levels in both humans and our mouse model. These observations suggest that Mcpip1^fl/fl^LysM^Cre^ mice meet the construct validity criteria by resembling molecular aspects of human liver inflammation.

In this work, we asked the question whether there is a biological target other than the regulation of systemic inflammation whose modulation might contribute to the reduction of symptoms of liver inflammation. We have found that modulation of LSEC activation may be an alternative and effective therapeutic strategy to counteract liver inflammation. This was achieved by treating Mcpip1^fl/fl^LysM^Cre^ mice with diosmetin. Diosmetin is a biologically active aglycone form of diosmin – a compound that is used in the clinic as a supportive therapy of chronic venous insufficiency and hemorrhoids^34,35^. We found that diosmetin treatment significantly decreased LSEC activation, improved its function by restoring cell surface glycocalyx, and, markedly reduced liver inflammation in Mcpip1^fl/fl^LysM^Cre^ mice. These observations meet predictive validity criteria, because diosmetin and its glycosylated form diosmin have also clinically proven anti-inflammatory vasoactive properties in humans.

While diosmetin’s hepatoprotective properties were previously suggested^59^, our study unveils the molecular mechanisms underlying its anti-inflammatory effect on the liver. Diosmetin inhibits TNFα-dependent activation of endothelial cells, by blocking the production of the NF-κB-p50 subunit, which is transcriptionally active form of NF-κB transcription factor^60^ involved in liver pathophysiology^61^. This was followed by a significant reduction in the expression levels of CAMs and increased levels of glycocalyx. Our observations are consistent with other studies reporting that a peptide designed to bind to NF-κB -p50 subunit could inhibit NF-κB activation and attenuated local acute inflammation^62^.

To sum up, in this study, we demonstrated several key findings:

1. In humans, reduced Mcpip1 expression in PBMC correlated with elevated plasma TNFα levels and LSEC dysfunction.
2. Mcpip1 deletion in myeloid cells in mice mirrored the molecular changes observed in human samples, including elevated TNFα and LSEC dysfunction.
3. Diosmetin treatment effectively blocked LSEC pro-inflammatory activation, preventing liver inflammation in Mcpip1^fl/fl^LysM^Cre^ mice despite ongoing systemic inflammation, suggesting its potential as an anti-inflammatory therapy of the liver in humans.

## Supporting information

Supplemental Metods

## Acknowledgements

This project was financially supported by research grant 2017/27/B/NZ5/01440 from National Science Centre, Poland.

## Data availability

All data files from the RNA-seq experiment are freely available in the BioProject collection under the accession number: PRJNA965950 (https://www.ncbi.nlm.nih.gov/bioproject/?term=PRJNA965950).

## Authors contributions

DŻ, JK, JJ– conceptualization and study design. DŻ, NP, KSz, KT, EK, EP, JK – experiments, procedures, data acquisition and analysis. P Major, P Małczak, DR, AB – clinical sample and data collection, clinical sample procedures. DŻ, JK, KSz, EP manuscript writing; DŻ, JK, JJ, SCh manuscript revision, DŻ manuscript editing. All authors approved final version of the manuscript.

## Abbreviations

ACTB: β-actin
AIH: autoimmune hepatitis
ALT: alanine transaminase
APTT: Activated Partial Thromboplastin Time
AST: aspartate transaminase
AU/px^2^: Auxiliary Units per squared pixel
BMI: Body mass index
CAMs: cell adhesion molecules
CHI3-L1: Chitinase 3 Like 1
CRP: C-reactive protein
CXCL16: Ligand chemokin 16
DEGs: differentially expressed genes
GGTP: gamma-glutamyl transpeptidase
Glx: glycocalyx
HBV-hepatitis B virus HCV: hepatitis C virus
HDL: high-density lipoprotein
HIV: human immunodeficiency virus
HUVEC: human umbilical vein endothelial cell
H&E: Hematoxylin and Eosin staining
ICAM-1: intercellular Adhesion Molecule 1
INR: international normalized ratio of protrombin time test
IL-6Rα: Interleukin-6 receptor-alpha
IκBα: nuclear factor of kappa light polypeptide gene enhancer in B-cells inhibitor alpha
LDL: low-density lipoprotein
LSEC: liver sinusoidal endothelial cell
MCP-5: Monocyte Chemotactic Protein-5
MCP-2: Monocyte Chemotactic Protein-2
Mcpip1: monocyte chemoattractant protein induced protein-1
Mcpip1^fl/fl^: control mice
Mcpip1^fl/flfl^LysM^Cre^: mice with Mcpip1 knock-out in myeloid cells
MIP-1α: macrophage inflammatory protein-1-alpha
MIP-3α: macrophage inflammatory protein-3-alpha
MIP-3β: macrophage inflammatory protein-3-beta
NAFLD: nonalcoholic fatty liver disease
NASH: steatohepatitis
NDE: number of DEGs in the pathway
NF-κB: nuclear factor kappa-light-chain-enhancer of activated B cells
PBMC: peripheral blood mononuclear cells
pNDE: hypergeometric probability
pPERT: bootstrap probability associated to tA,
p-Size: the number of genes in the pathway
p-val: nominal p-value
q-val: FDR adjusted p-value
Selp: selectin P
SDC1: syndecan-1
tA: perturbation score value
TNFα: tumor necrosis factor alpha
TNFRI: TNFα receptor type I
WBC: white blood cells
VCAM-1: vascular cell adhesion molecule 1
VEGFRII: Vascular endothelial growth factor receptor 2
vWF: von Willebrandt factor.

---

**Supplementary table 1 Demographical and biochemical data from patients with high and low plasma levels of TNFα.**

Demographic and blood biochemical analysis data are presented as mean ± SEM. p-values for all biochemical parameters were calculated by Mann-Whitney test. Abbreviations: ALT - alanine transaminase, APTT - Activated Partial Thromboplastin Time, AST - aspartate transaminase, BMI - Body mass index, CRP - C-reactive protein, GGTP - gamma-glutamyl transpeptidase, HDL - high- density lipoprotein, INR - international normalized ratio of protrombin time test, LDL - low-density lipoprotein, WBC - white blood cells.

**Fig. S1 TNFα is the only cytokine elevated in Mcpip1^fl/fl^LysM^Cre^ mice that mediates the pro-inflammatory activation of endothelial cells *in vitro*.**

HUVEC endothelial cells were stimulated *in vitro* with six cytokines (TNFα, MCP-2, MCP-5, MIP-3α, MIP-3β and MIP-1α) that were found to be elevated in Mcpip1^fl/fl^LysM^Cre^ mice to test which one has a potential to activate endothelial cells. Protein expression levels of **(A)** ICAM1 (n=6), **(B)** VCAM1 (n=6), **(C)** SELP (n=6) and **(D)** SDC1 (n=6) were tested in HUVEC cells stimulated by three different concentrations of each cytokine (see Supplementary file 1). Blue triangles represent increasing concentrations of each cytokine Non-stimulated cells were used as a control. Data are presented as mean ± SEM in the bar graphs. P-values were calculated by using one-way ANOVA followed by FDR post hoc test. Representative western blots of ICAM1, VCAM1, SELP, SDC1 and ACTB from cells stimulated with three different concentrations of: **(E)** TNFα. **(F)** MCP-2. **(G)** MCP-5. **(H)** MIP-1α. **(I)** MIP-3β. **(J)** MIP-3α. Abbreviations: ACTB – β-actin, ICAM1 - intercellular Adhesion Molecule 1, MCP-2 Monocyte Chemotactic Protein-2, MCP-5 - Monocyte Chemotactic Protein-5, MIP-1 α - macrophage inflammatory protein-1-alpha, MIP-3β - macrophage inflammatory protein-3-beta, MIP-3α - macrophage inflammatory protein-3-alpha, SELP – selectin P, SDC1 – Syndecan 1, TNFα – tumor necrosis factor alpha, VCAM1 - vascular cell adhesion molecule 1.

**Fig. S2 Diosmetin blocks the production of NF-κB-p50 subunit and decreases TNFα-dependent pro-inflammatory activation of endothelial cells *in vitro*.**

Protein expression levels of **(A)** ICAM1 (n=4), **b)** VCAM1 (n=4), **(C)** SELP (n=4), **(D)** SDC1 (n=4), **(E)** NF-κB inhibitory protein IκBα (n=4), **(F)** NF-κB-p105 (n=4), and **(G)** NFκB-p50 subunit (n=4) were tested in in HUVEC endothelial cells co-stimulated with diosmetin (in three different concentrations) and TNFα (in two different concentrations). Non-stimulated cells were used as a control. Data are presented as mean ± SEM in the bar graphs. P-values were calculated by using one-way ANOVA followed by FDR post hoc test. **(H)** Representative western blots of ICAM1, VCAM1, SELP, SDC1 and ACTB from cells co-stimulated with TNFα and diosmetin. **(I)** Representative western blots of IκBα, NF- κB-p105, NFκB-p50 and ACTB from cells co-stimulated with TNFα and diosmetin. Abbreviations: ACTB – β-actin, DIOS – diosmetin, ICAM1 - intercellular Adhesion Molecule 1, IκBα - NF-κB inhibitory protein, NF-κB - nuclear factor kappa-light-chain-enhancer of activated B cells, NF-κB-p105 - NF-κB precursor protein 105, SELP – selectin P, SDC1 – Syndecan 1, TNFα – tumor necrosis factor alpha, VCAM1 - vascular cell adhesion molecule 1.

