## Supplemental Metods for "Diosmetin alleviates liver inflammation by improving liver sinusoidal endothelial cell dysfunction"

### Supplementary file 1

#### Diosmetin alleviates liver inflammation by improving liver sinusoidal endothelial cell dysfunction.

Dariusz Żurawek[1,2,#], Natalia Pydyn[1], Piotr Major[3], Krzysztof Szade[4], Katarzyna Trzos[1], Edyta Kuś[5], Ewelina Pośpiech[6], Piotr Małczak[3], Dorota Radkowiak[3], Andrzej Budzyński[3], Stefan Chłopicki[5], Jolanta Jura[1],Jerzy Kotlinowski[1,#]

#### Complete list of PCR primers, cytokines, primary and secondary antibodies (and their dilutions) used in the analysis:

##### Table S1. List of mouse genotyping primers.

| Primer sequence | Type |
|---|---|
| GCCTTCCTGATCCTATTGGAG | Wild-type |
| GAGATGGCGCAGCGCAATTAAT | Knock-out |
| GCCTCTTGTCACTCCCTCCTCC | common |
| TTACAGTCGGCCAGGCTGAC | LysMCre $^{tg/+}$ wild-type |
| CCCAGAAATGCCAGATTACG | LysMCre $^{tg/+}$ mutant |
| CTTGGGCTGCCAGAATTTCTC | LysMCre $^{tg/+}$ common |

##### Table S2. List of primers used for RT-PCR.

| Gene | Forward (5' -> 3') | Reverse (5' -> 3') | NCBI no. |
|---|---|---|---|
| **Ef2** | GACATCACCAAGGGTGTGCAG | TTCAGCACACTGGCATAGAGGC | NM_001961 |
| **Sdc1** | ACAGAGCCTAACGCAGAGGAA | GAGCGAGGGCTGTAGGTTTC | NM_011519.2 |
| **Icam1** | GGAGCCAATTTCTCATGCCG | GTGTCGAGCTTTGGGATGGT | NM_010493.2 |
| **Vcam1** | CTAATTCATGGTAGAATGGCTA | TGAAGTCGCATTTAAATCAGGT | NM_011693.3 |
| **vWf** | CTCCAGCCACATTCCATACC | GAGATGGGCGTAAGAAGCAA | NM_011708.4 |
| **Selp** | ACAATGTCCTGCCAACACCA | GTCCATGTGCAATTCGGAGC | NM_011347.2 |

##### Table S3. Antibodies and reagents used for flow cytometry. All antibodies were used at the 1:50 dilution.

| Antigen | Dye | Clone | Company |
|---|---|---|---|
| CD3 | AlexaFluor700 | 17A2 | BioLegend, USA (Cat# RUO 100216) |
| CD11b | APC-Cy7 | M1/70 | BD Biosciences, USA (Cat# RUO557657) |
| CD16/CD32 | - | 2.4G2 | BD Biosciences, USA (Cat# RUO553141) |
| CD45 | Pacific Orange | 30F-11 | ThermoFisher, USA (Cat# MCD4530) |
| CD45R (B220) | PE | RA3-6B2 | BD Biosciences, USA (Cat# RUO553089) |
| Ly6C | PerCP-Cy5.5 | HK1.4 | eBioscience, USA (Cat# 45-5932-82) |
| Ly6G | FITC | 1A8 | BD Biosciences, USA (Cat# RUO551460) |
| I-A/I-E (MHC II) | PE-Cy7 | M5/114.15.2 | eBioscience, USA (Cat# 25-5321-82) |
| Ly71 (F4/80) | APC | BM8 | eBioscience, USa (Cat# 17-4801-82) |
| - | DAPI | - | MiliporeSigma, USA (Cat# 28718-90-3) |

#### Table S4. Antibodies used for immunohistochemistry.

| Antigen | Dye | Dilution | Company |
|---|---|---|---|
| Selp | - | 10ug/ml | R & D Systems, USA (Cat# AF737) |
| Vcam-1 | - | 1:500 | abcam, UK (Cat# ab134047) |
| Icam-1 | - | 1:250 | abcam, UK (Cat# ab108361) |
| Goat IgG | Alexa Fluo488 | 1:1000 | Thermo Fisher, USA (Cat# A-11055) |
| Rabbit IgG | Alexa Fluo546 | 1:1000 | Thermo Fisher, USA (Cat# A-11035) |
| Rabbit IgG | Alexa Fluo488 | 1:1000 | Thermo Fisher, USA (Cat# A-11008) |

#### Table S5. Antibodies used for immunoblotting.

| Antigen | Dilution | Company |
|---|---|---|
| ICAM-1 | 1:1000 | abcam, UK (Cat# ab109361) |
| VCAM-1 | 1:1000 | abcam, UK (Cat# ab134047) |
| SELP | 1:1000 | Thermo Fisher, USA (Cat# PA5-88593) |
| SDC1 | 1:2000 | abcam, UK (Cat# ab128936) |
| ACTB | 1:2000 | Merck Milipore, Germany) |
| p50/p105 NFκB | 1:1000 | Cell Signaling, Netherlands (Cat#30355) |
| IκBα | 1:1000 | Cell Signaling, Netherlands (Cat#4814) |
| Mcpip1 | 1:1000 | Abcam, UK (Cat# ab252879) |
| Rabbit IgG | 1:20 000 | MilliporeSigma, Germany (Cat# SAB3700934) |
| Mouse IgG | 1:10 000 | BD Biosciences, USA (Cat# 554002) |

#### Table S6. List of cytokines and their concentrations used in vitro to stimulate HUVEC cells.

| Cytokine | Dilution 1 (corresponding to average plasma concentration in wild-type mice) | Dilution 2 (corresponding to average plasma concentration in Mcpip1$^{fl/fl}$LysM$^{Cre}$ mice) | Dilution 3 (corresponding to concentration recommended by the manufacturer) | Company |
|---|---|---|---|---|
| TNFα | 2pg/ml | 8pg/ml | 10ng/ml | Merck Milipore, Germany(Cat# 654205-M) |
| MCP-2 | 0.1ng/ml | 1.2ng/ml | 10ng/ml | R & D Systems, USA (Cat# 281-CP) |
| MCP-5 | 0.1pg/ml | 0.3pg/ml | 10ng/ml | R & D Systems, USA (Cat# 428-P5) |
| MIP-1α | 1pg/ml | 5pg/ml | 10ng/ml | R & D Systems, USA (Cat# 270-LD) |
| MIP-3β | 0.2pg/ml | 2pg/ml | 10ng/ml | abcam, UK (Cat# ab9832) |
| MIP-3α | 0.2pg/ml | 2pg/ml | 10ng/ml | abcam, UK (Cat# ab256086) |
