## Supplemental Table for "Diosmetin alleviates liver inflammation by improving liver sinusoidal endothelial cell dysfunction"

**Demographical and biochemical data from patients with high and low plasma levels of TNFα**

| | Control | High TNFα | p-value |
|---|---|---|---|
| n | 18 | 24 | n/a |
| Sex | 12F/6M | 20F/4M | n/a |
| Ethnicity | Caucasian 100% | Caucasian 100% | n/a |
| Age | 45,56 ± 2,772 | 45,84 ± 2,586 | 0,9417 |
| BMI | 45,98 ± 2,813 | 40,69 ± 2,048 | 0,1266 |
| CRP [mg/L] | 37,58 ± 5,165 | 51,23 ± 13,23 | 0,6188 |
| WBC [$10^3$/ul] | 11,51 ± 0,6414 | 11,74 ± 0,5699 | 0,7903 |
| Neutrophils [$10^3$/ul] | 8,9 ± 0,524 | 8,9 ± 0,556 | 0,9008 |
| Lymphocytes [$10^3$/ul] | 1,66 ± 0,14 | 1,51 ± 0,13 | 0,4097 |
| Monocytes [$10^3$/ul] | 0,89 ± 0,06 | 0,94 ± 0,07 | 0,529 |
| Red blood cells [$10^6$/ul] | 4,47 ± 0,073 | 4,51 ± 0,097 | 0,7379 |
| Hemoglobin [g/dL] | 13,02 ± 0,28 | 13,20 ± 0,28 | 0,6686 |
| Platelets [$10^3$/ul] | 243,8 ± 13,91 | 230 ± 9,062 | 0,3958 |
| INR | 1,009 ± 0,011 | 0,984 ± 0,011 | 0,1064 |
| APTT [sec] | 27,06 ± 0,58 | 27,71 ± 0,68 | 0,4838 |
| Hematocrit [%] | 39,14 ± 0,71 | 38,9 ± 0,76 | 0,8238 |
| AST [IU/L] | 124,2 ± 40,91 | 103,6 ± 29,02 | 0,6898 |
| ALT [IU/L] | 191 ± 69,55 | 164,3 ± 53,42 | 0,2976 |
| GGTP [IU/L] | 52,06 ± 9,734 | 52,19 ± 15,11 | 0,3981 |
| Bilirubin [umol/L] | 11,3 ± 1,043 | 11,92 ± 1,614 | 0,6593 |
| Sodium [mmol/L] | 140,4 ± 0,6 | 140,2 ± 0,52 | 0,7782 |
| Potasium [mmol/L] | 4,15 ± 0,08 | 4,19 ± 0,07 | 0,7379 |
| Iron [mmol/L] | 9,74 ± 0,83 | 8,89 ± 0,78 | 0,4606 |
| Urea [mmol/L] | 4,46 ± 0,34 | 4,89 ± 0,79 | 0,8984 |
| Creatinine [mmol/L] | 76,67 ± 3,95 | 81,39 ± 5,48 | 0,7501 |
| Glucose [mmol/L] | 6,44 ± 0,33 | 7,41 ± 0,58 | 0,2601 |
| Albumin [mg/ml] | 41,14 ± 0,76 | 41,72 ± 0,87 | 0,6233 |
| Cholesterol [mmol/L] | 4,4 ± 0,19 | 4,26 ± 0,17 | 0,5951 |
| HDL [mmol/L] | 1,2 ± 0,05 | 1,24 ± 0,08 | 0,672 |
| LDL [mmol/L] | 2,63 ± 0,19 | 2,51 ± 0,16 | 0,6101 |
| Triglycerides [mmol/L] | 1,58 ± 0,15 | 1,47 ± 0,12 | 0,5526 |

Values in the table refer to the mean ± SEM.
p-values were calculated by non-parametric Mann-Whitney test

Abbreviations: ALT - alanine transaminase, APTT - Activated Partial Thromboplastin Time, AST - aspartate transaminase, BMI - Body mass index, CRP - C-reactive protein, GGTP - gamma-glutamyl transpeptidase, HDL - high-density lipoprotein, INR - international normalized ratio of protrombin time test, LDL - low-density lipoprotein, WBC - white blood cells
