## Supplementary figures and images for "Diosmetin alleviates liver inflammation by improving liver sinusoidal endothelial cell dysfunction"

### Supplemental Figure 1

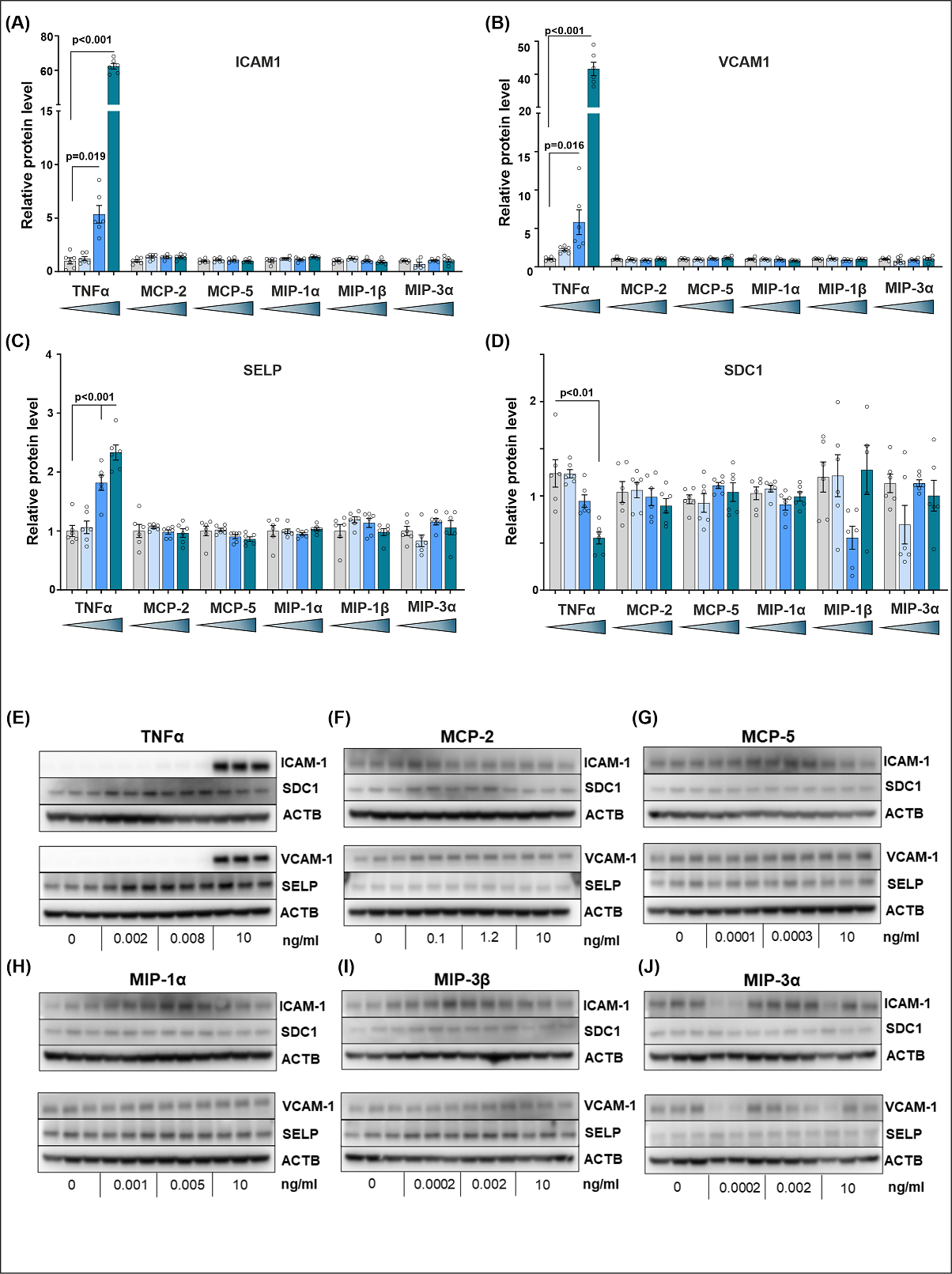

### Supplemental Figure 2

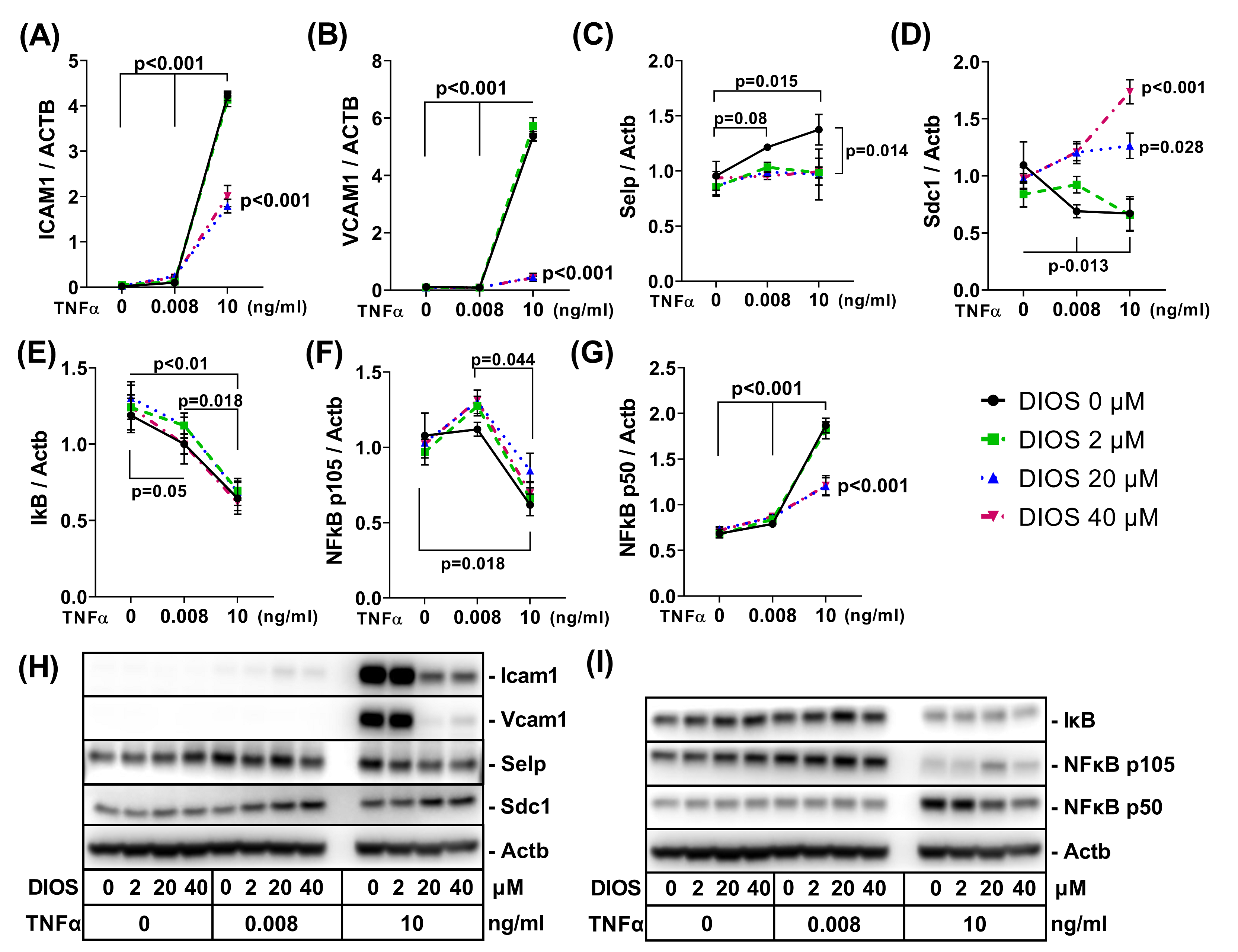
